# PIP_2_ and Ca^2+^ regulation of TMEM16A currents in excised inside-out patches

**DOI:** 10.1101/2022.08.30.505925

**Authors:** Maiwase Tembo, Crystal Lara-Santos, Joel C. Rosenbaum, Anne E. Carlson

## Abstract

The Ca^2+^ activated Cl^−^ channel formed by transmembrane member 16A (TMEM16A) is broadly expressed and regulates diverse processes. In addition to Ca^2+^, TMEM16A channels require the acidic phospholipid phosphatidylinositol 4,5-bisphosphate (PI(4,5)P2) to open. Like other channels regulated by PI(4,5)P_2_, TMEM16A-conducted currents recorded in excised patches slowly decay overtime. Here we assessed how intracellular Ca^2+^ alters the rate of this current rundown, using the channels endogenously expressed in oocytes from the African clawed frog, *Xenopus laevis*. We found that in excised, inside-out patches, the concentration of applied Ca^2+^ alters the rate of rundown, with high Ca^2+^ concentrations speeding rundown by activating membrane associated phospholipase C (PLC). Together, these results clarify our understanding of how Ca^2+^ regulates both TMEM16A directly, and targets PLC to regulate the membrane PI(4,5)P_2_ content.

## 1 Introduction

The Ca^2+^ -activated Cl^−^ channel, TMEM16A, is broadly expressed and regulates diverse physiologic processes ranging from mucosal secretion (Cabrita et al., 2021; Huang et al., 2012), transduction of sensory signals (Chen et al., 2021; Hernandez-Clavijo et al., 2021), smooth muscle tone (Bulley et al., 2012; Danielsson et al., 2020; Wang et al., 2021), and fertilization (Wozniak, Phelps, Tembo, Lee & Carlson, 2018; Wozniak, Tembo, Phelps, Lee & Carlson, 2018). Because TMEM16A channels regulate these important yet varied processes, it could be targeted by novel therapeutics as treatments for chronic diseases such as hypertension.

TMEM16A channels are homodimers, with each subunit comprising ten transmembrane domains, large intracellular N- and C-terminal domains, a membrane-embedded Ca^2+^ binding site, and an independently operating Cl^−^ conducting pore (Al-Hosni, Ilkan, Agostinelli & Tammaro, 2022; Hawn, Akin, Hartzell, Greenwood & Leblanc, 2021; Shi et al., 2020). Five conserved amino acids in transmembrane domains 6-8 form the Ca^2+^ binding site, with each site capable of accommodating two Ca^2+^ ions (Brunner, Lim, Schenck, Duerst & Dutzler, 2014; Dang et al., 2017; Paulino, Kalienkova, Lam, Neldner & Dutzler, 2017). The channels gates differentially depending on whether one or two Ca^2+^ ions are bound (Peters et al., 2018). Moreover, other divalent cations in addition to Ca^2+^ are capable of opening TMEM16A channels (Yuan et al., 2013).

In addition to divalent cations, TMEM16A channels also require the acidic phospholipid phosphatidylinositol 4,5-bisphosphate (PI(4,5)P2) to open (De Jesus-Perez et al., 2017; Le, Jia, Chen & Yang, 2019; Ta, Acheson, Rorsman, Jongkind & Tammaro, 2017; Tembo et al., 2022; Tembo, Wozniak, Bainbridge & Carlson, 2019; Yu, Jiang, Cui, Tajkhorshid & Hartzell, 2019). A hallmark of channels potentiated by PI(4,5)P2 is that their currents rundown when recorded in excised patches (Suh & Hille, 2008). Indeed, TMEM16A conducted currents decay in excised patches, even in the presence of continuously applied intracellular Ca^2+^ (Tembo et al., 2022; Tembo, Wozniak, Bainbridge & Carlson, 2019). How PI(4,5)P2 potentiates TMEM16A conductance is unknown. One hypothesis states that PI(4,5)P2 stabilizes the open state of permeation pore of TMEM16A channels by binding to a separate regulatory module (Le, Jia, Chen & Yang, 2019), while another suggests that PI(4,5)P2 opens the pore by direct interaction with the outside of a pore-lining helix (Yu, Jiang, Cui, Tajkhorshid & Hartzell, 2019).

The full length TMEM16A isoform requires the phosphate at the 4’ position of PI(4,5)P2 in order to activate these channels (Ko & Suh, 2021; Tembo et al., 2022). Whereas the full-length channel is widely expressed in mammals, a TMEM16A splice variant expressed in the brain and skeletal muscle lacks four residues (EAVK) that reside in a linker connecting transmembrane domains 2 and 3; for this variant, the phosphate position does not matter for the phospholipid to potentiate channel activity (Ko & Suh, 2021; Le, Jia, Chen & Yang, 2019).

Notably, the lysine of the EAVK motif potentially interacts with the 4’ phosphate of PI(4,5)P_2_. Here we explored the relationship between intracellular Ca^2+^ on both the regulation of TMEM16A and the rate of current rundown in excised inside-out patches. Using electrophysiology recordings made on oocytes from the African clawed frog, *Xenopus laevis*, a cell that abundantly expresses TMEM16A channels, we found that millimolar Ca^2+^ application both activated TMEM16A conducted currents, but that these currents quickly decayed in excised patches. By contrast, the current rundown was significantly slower when the lower concentration of Ca^2+^ was applied at 500 µM, or when the channel was activated by 2 mM of other divalent cations including Ni^2+^ or Ba^2+^. Together these data reveal that 2 mM Ca^2+^ activates TMEM16A as well as phospholipase C (PLC) enzymes associated with the patch to speed PI(4,5)P_2_ depletion.

## 2 Methods

### 2.1 Animals

Animal procedures used were approved by the Animal Care and Use Committee at the University of Pittsburgh and are consistent with the accepted standards of humane animal care. *Xenopus laevis* adult, oocyte-positive females were obtained commercially (NASCO, Fort Atkinson, WI or Xenopus 1, Dexter MI) and housed at 18-20 °C with a 12/12-hour light/dark cycle.

### 2.2 Oocyte Collection

*X. laevis* females were anesthetized by immersion in 1.0 g/L tricaine pH 7.4 for 30 min before oocytes were collected. The ovarian sacs were surgically obtained and manually pulled apart using blunt forceps. Oocytes were treated with a 90-min incubation in 1 mg/ml collagenase in the ND96 solution, then repeatedly rinsed in OR2 to remove collagenase. Healthy oocytes were stored at 12-14 °C in pyruvate- and gentamycin-supplemented ND96 solution.

### 2.3 Solutions

Inside-out patch-clamp recordings were conducted in a HEPES-buffered saline solution made as follows (in mM): 130 NaCl and 3 HEPES, pH 7.2, and filtered using a sterile, 0.2 µm polystyrene filter (Tembo, Wozniak, Bainbridge & Carlson, 2019). The HEPES-buffered saline solution was supplemented with 0.2 mM EGTA for Ca^2+^-free recordings. For recordings made with intracellular Ca^2+^, the HEPES-buffered saline solution was supplemented with 2 mM, or 500 µM CaCl_2_ with indicated reagents. Total Ca^2+^ concentrations are reported throughout the manuscript. Free Ca^2+^ concentrations were calculated using Maxchelator (Bers, Patton & Nuccitelli, 2010) and are 1.8 mM Ca^2+^ (with 2 mM total Ca^2+^) and 300 µM Ca^2+^ (with 500 µM total Ca^2+^). The diC8-PI(4,5)P2 analog was added to the 2 mM Ca^2+^-containing HEPES-buffered saline solution and applied as indicated.

The oocyte wash solution, called Oocyte Ringers 2, and storage solution, ND96, were made as follows. Oocyte Ringers 2 (OR2) (in mM): 82.5 NaCl, 2.5 KCl, 1 MgCl2, and 5 mM HEPES, pH 7.2. ND96 (in mM): OR2 supplemented with 5 mM sodium pyruvate 100 mg/L gentamycin, pH 7.6, and 0.2 µm polystyrene filtered.

### 2.4 Patch-Clamp Recordings

Patch-clamp recordings were conducted on *X. laevis* oocytes following the manual removal of the vitelline membrane. TMEM16A current recordings were made in the inside-out configuration of the patch-clamp technique using an EPC-10 USB patch-clamp amplifier (HEKA Elektronic). The Patchmaster software (HEKA Elektronic) was used for data acquisition. Briefly, following the establishment of a gigaseal (greater than 1 GΩ), patches were excised in HEPES-buffered saline solution lacking both EGTA and added calcium. Following excision in this EGTA-free, HEPES-buffered saline, the patch resistances typically decreased to 20-200 MΩ, but returned to >1 GΩ with EGTA application. Data were collected at a rate of 10 kHz. Glass pipettes were pulled from borosilicate glass (OD 1.5 mm, ID 0.86 mm; Warner Instruments) and were fire polished using a Narshige microforge. Pipettes resistance ranged from 0.4-1.5 MΩ. All experiments were initiated within 10 s of patch excision. A VC-8 fast perfusion system (Warner Instruments) was used for solution application.

Patch-clamp data were analyzed with Excel (Microsoft) and IGOR Pro (Wavemetrics) with Patchers Power Tools. Currents were processed such that peak currents were normalized to 1. To calculate the rate of rundown, plots of normalized current at -60 mV versus time were fit with the single exponential equation:

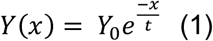

where Y_0_, x, t, and Y(x) represent the initial current, time, rate of current rundown, and current at time x (Tembo, Wozniak, Bainbridge & Carlson, 2019). Remaining current at 100, 150, and 200 s was assessed by determining the proportion of maximum current at each of these time points (relative to the beginning of the experiment and included 10 s of EGTA-containing, no Ca^2+^ added solution application).

To calculate the current recovered following application of the synthetic PIP_2_ analog, diC8-PIP_2_, the fold change in current recovered was calculated by dividing the peak current after diC8-PIP_2_ addition by the baseline current. The peak current was defined as the highest current obtained after diC8-PIP_2_ addition. The baseline current was defined as the current observed at the point of diC8-PIP_2_ addition. The equation used was:

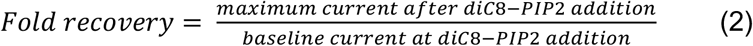

A fold recovery of 1 relates to unchanged current, and a recovery > 1 indicates that diC8-PIP_2_ application increased the current.

## 3 Results

### 3.1 TMEM16A currents decayed in excised patches and were recovered by PI(4,5)P_2_

Using *X. laevis* oocytes, we recorded Ca^2+^-activated TMEM16A-conducted currents using the excised-inside-out configuration of the patch clamp technique. Briefly, currents were recorded during 150 ms steps to -60 and +60 mV before and during application of 2 mM Ca^2+^. As we previously reported, application of 2 mM Ca^2+^ evoked robust TMEM16A-conducted currents immediately following patch excision (Tembo et al., 2022; Tembo, Wozniak, Bainbridge & Carlson, 2019). Figure 1A depicts currents recorded during steps to -60 and +60 mV at indicated time points (10-180 s) following 2 mM Ca^2+^ addition. Despite the continued application of intracellular Ca^2+^, these currents decayed over time (Figure 1). By fitting single exponential functions (Equation 1) to plots of normalized currents recorded during steps to -60 mV versus time (Figure 1B), we found that currents decayed with an average tau (τ) of 90.1 ± 8.3 s (N = 18).

**Figure 1:**
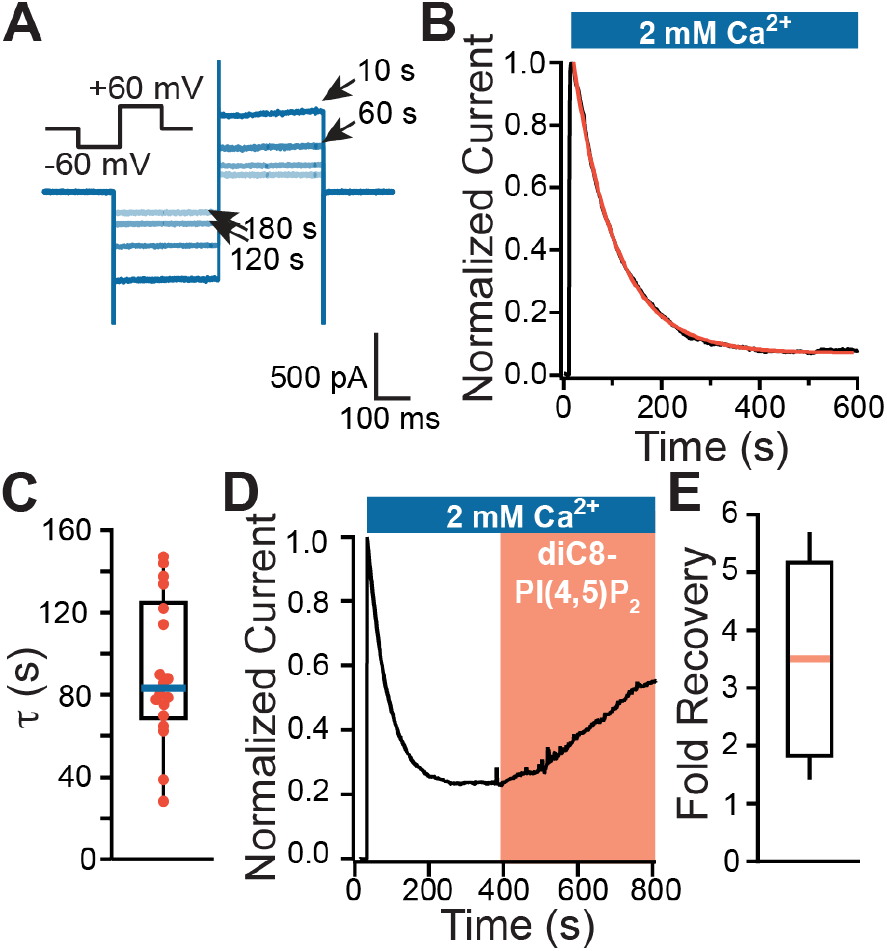
TMEM16A currents rundown in excised inside-out patches. Excised inside-out patches clamp recordings were obtained from macropatches made on *X. laevis* oocytes. (A) Example currents recording during 150-ms steps to -60 and +60 mV at the given times after patch excision. (B) Normalized plot of currents vs time after patch excision. 2 mM Ca^2+^ was applied at 10 s, as denoted by the blue bar, and currents were fitted with a single exponential (red line). (C) Box plot distribution of rate of current decay (τ) obtained from fits with single exponentials (N = 18). The box represents 25-75% of the data distribution, the whiskers indicate 0% to 100% of the data, and the center line represents the median. (D) Example plot of relative current recorded at -60 mV vs time of an inside-out patch before and during application of 100 µM diC8-PI(4,5)P_2_ (N = 6). The diC8-PI(4,5)P_2_ analog was applied following rundown, once currents reached a steady state. (E) Box plot distribution of the fold-recovery of current after 100 µM diC8-PIP_2_ application.

Current rundown is a characteristic of channels regulated by the phospholipid PI(4,5)P_2_ when recorded using the inside-out configuration of the patch-clamp technique (Suh & Hille, 2008). PI(4,5)P_2_ is depleted in excised patches; indeed, we found that application of 100 µM of the soluble dioctanoyl-phosphatidyl 4,5-bisphosphate (diC8-PI(4,5)P_2_) recovered TMEM16A currents. Figure 1D shows an example plot of TMEM16A-conducted currents recorded at -60 mV versus time, before and during application of 100 µM diC8-PI(4,5)P_2_. By calculating the change in current with diC8-PI(4,5)P_2_ application (Equation 2), we observed that diC8-PI(4,5)P_2_ recovered an average of 3.5 ± 0.7-fold current (N=6, Figure 1D&E).

### 3.2 TMEM16A current decay was slower with other divalent cations

To explore TMEM16A regulation by other divalent cations, we applied Ni^2+^ or Ba^2+^ to inside-out patches excised from *X. laevis* oocytes (Yuan et al., 2013) (Figure 2A). We found that when applied at 2 mM, both Ni^2+^ and Ba^2+^ activated TMEM16A channels (Figures 2B&C), however, the current decay differed substantially from that in 2 mM Ca^2+^. For example, plots of the relative current in the presence of 2 mM Ni^2+^ were not well fit with single exponentials (Figure 2B), thus, we assessed the proportion of maximum current at 100, 150, and 200 s (Table 1). When activated with 2 mM Ni^2+^, the TMEM16A-conducted current at 100 s only decayed to 0.80 ± 0.05 proportion of maximum current (N=10). By contrast, the currents decayed significantly more in 2 mM Ca^2+^, with only 0.45 ± 0.05 remaining at 100 s (N=18) (*P*<0.01, T-test, Figure 2B). Similarly, the TMEM16A current decayed more slowly in 2 mM Ba^2+^. Plots of relative current versus time were well fit with single exponentials and these Ba^2+^ activated TMEM16A conducted currents decayed with an averaged τ of 361.2 ± 121.0 s (N=5, Figure 2E), substantially slower than in 2 mM Ca^2+^ (90.1 ± 8.3 s, *P*<0.01, T-test). At 100 s following excision, 0.74 ± 0.08 current remained in Ba^2+^, which was not significantly different from the current remaining at 100 s in Ni^2+^ (Figure 2B). Together these data establish that Ba^2+^ and Ni^2+^ activate TMEM16A, but even at millimolar concentrations, the rate of rundown in the presence of these divalent cations is substantially slower than in 2 mM Ca^2+^. Evidently, all three divalent cations activate TMEM16A, but Ca^2+^ additionally speeds rundown.

**Table 1:** Relative remaining current during rundown under different conditions. The mean ± S.E.M. for of the normalized currents at the indicated time, taken from excised inside-out patches.

| Treatment | N | 100 s | 150 s | 200 s |
| --- | --- | --- | --- | --- |
| 2 mM Ca<sup>2+</sup> | 18 | 0.45 ± 0.05 | 0.28 ± 0.04 | 0.19 ± 0.03 |
| 2 mM Ni<sup>2+</sup> | 10 | 0.80 ± 0.05 | 0.71 ± 0.08 | 0.62 ± 0.10 |
| 2 mM Ba<sup>2+</sup> | 5 | 0.74 ± 0.08 | 0.64 ± 0.10 | 0.55 ± 0.11 |
| 500 μM Ca<sup>2+</sup> | 11 | 0.67 ± 0.06 | 0.50 ± 0.08 | 0.39 ± 0.08 |
| 2 mM Ca<sup>2+</sup> and ATP | 7 | 0.70 ± 0.05 | 0.56 ± 0.07 | 0.45 ± 0.07 |
| 500 μM Ca<sup>2+</sup> and ATP | 8 | 0.83 ± 0.03 | 0.77 ± 0.03 | 0.72 ± 0.04 |
| 2 mM Ca<sup>2+</sup> and U73122 | 5 | 0.71 ± 0.09 | 0.58 ± 0.05 | 0.47 ± 0.10 |
| 2 mM Ca<sup>2+</sup> and PMSF | 6 | 0.75 ± 0.08 | 0.60 ± 0.08 | 0.43 ± 0.08 |

**Figure 2:**
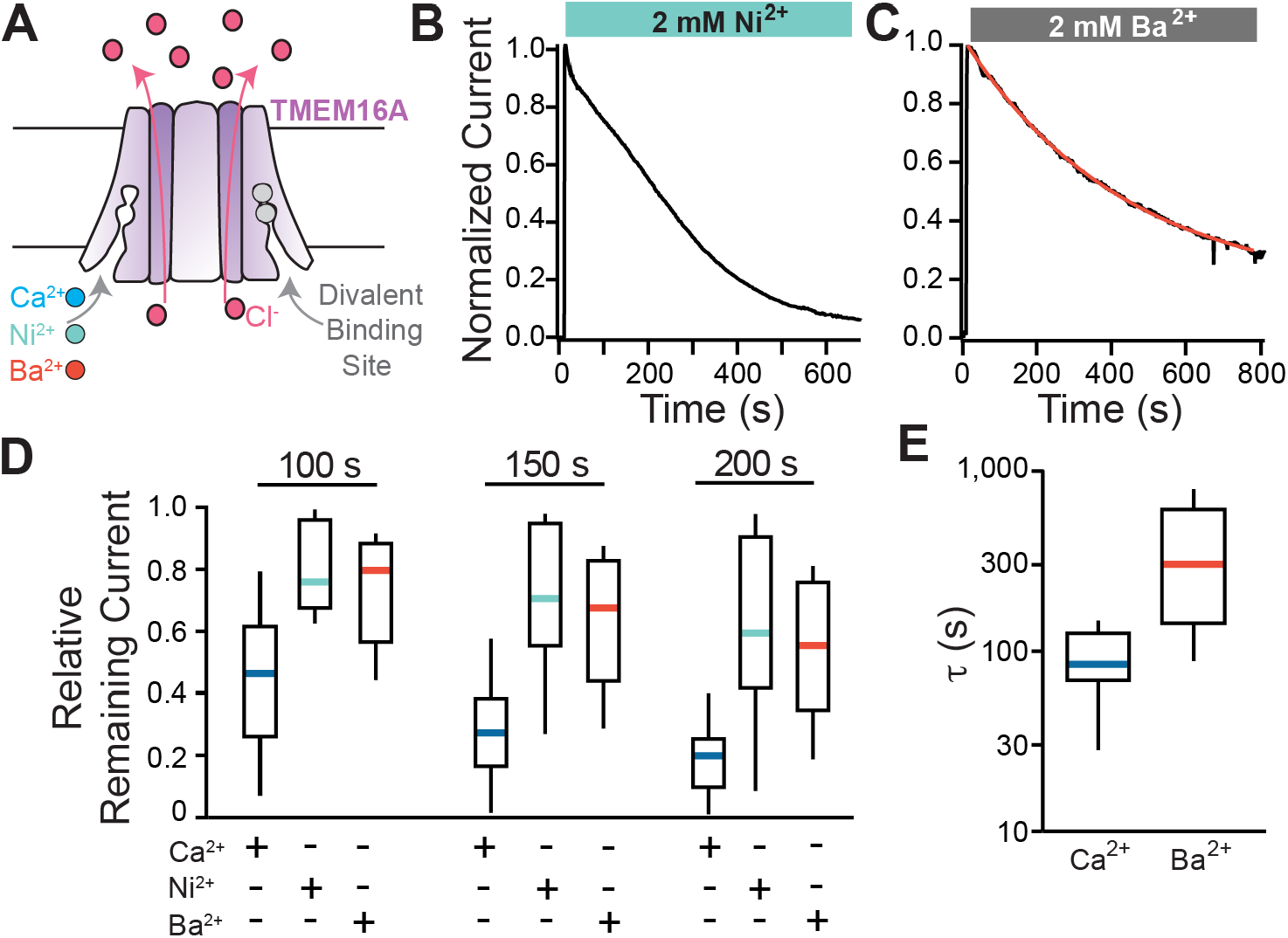
TMEM16A currents decayed more slowly with Ni^2+^ or Ba^2+^ activation. (A) Schematic of TMEM16A binding site for divalent cations Ca^2+^, Ni^2+^, or Ba^2+^. (B-C) Representative plot of normalized currents versus time before and during application of 2 mM Ni^2+^ (B) or 2 mM Ba^2+^ (C). Divalent cations were applied at 10 s. Plots of normalized current in Ba^2+^ vs time was fit with a single exponential (red line). (D) Box plot distribution of relative remaining current at three time points following patch excision: 100, 150, and 200 s (N=5-18). (E) Box plot distribution of rate of current decay (τ) obtained from fits with single exponentials (N = 5).

### 3.3 Applied Ca^2+^ concentration altered the rate of TMEM16A current rundown

We next quantified the rate of TMEM16A current rundown at the lower Ca^2+^ concentration of 500 µM. In eleven independent trials, 500 µM Ca^2+^ activated TMEM16A conducted currents and that these currents decayed over time (Figure 3A&B). By fitting single exponentials to plots of normalized current vs time, we observed slower rundown in 500 µM vs 2 mM Ca^2+^, with an averaged τ of 193.4 ± 28.4 s compared to 91.0 ± 8.3 s (*P*=<0.01, T-test) (Figure 3). These data indicate that the applied concentration of Ca^2+^ alters the rate of rundown and suggest that millimolar Ca^2+^ targets TMEM16A channels while also regulating the rate of PIP2 depletion.

**Figure 3:**
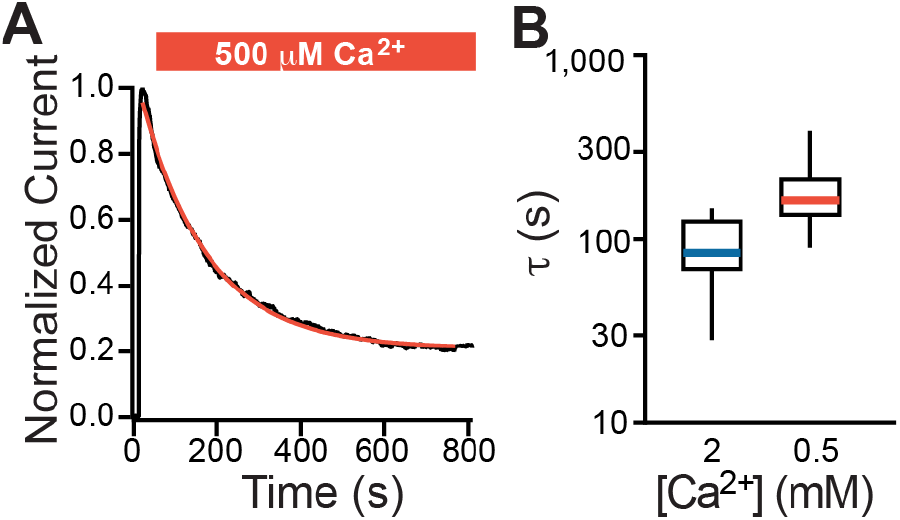
Slower rundown of TMEM16A currents at a lower Ca^2+^ concentration. (A) Representative plot of normalized current recorded at -60 mV vs time of TMEM16A conducted currents in the presence of 500 µM Ca^2+^ denoted by the red bar. Plot was fit with a single exponential (red line) (B) Box plot distribution of the rundown rate (τ) during application of either 2 mM or 500 µM Ca^2+^ (N = 11-18).

### 3.4 ATP application partially rescued rundown in 2 mM Ca^2+^

We reasoned that in addition to activating TMEM16A, 2 mM Ca^2+^ could target other membrane associated proteins to speed PI(4,5)P_2_ depletion in excised patches. Because the rundown of TMEM16A currents in excised patches is due to PI(4,5)P_2_ depletion (Tembo et al., 2022; Tembo, Wozniak, Bainbridge & Carlson, 2019), we explored a possible role for Ca^2+^ in the regulation of PI(4,5)P_2_ homeostasis in excised patches. In living cells, phosphatases and kinases work together to maintain membrane PI(4,5)P_2_ content; in excised patches, membrane-anchored kinases lack access to the ATP required to fuel phosphorylation and regeneration of PI(4,5)P_2_ (Figure 4A) (Hilgemann & Ball, 1996; Suh & Hille, 2008). Continued and unopposed activity of phosphatases in excised patches therefore leads to the net dephosphorylation of PIP_2_ (Figure 4A). We have previously reported that application of 2 mM Ca^2+^ with ATP slowed rundown of TMEM16A conducted current recorded from excised inside out patches (Tembo, Wozniak, Bainbridge & Carlson, 2019). Here we compared how ATP application altered rundown in patches treated with 2 mM or 500 µM Ca^2+^.

**Figure 4:**
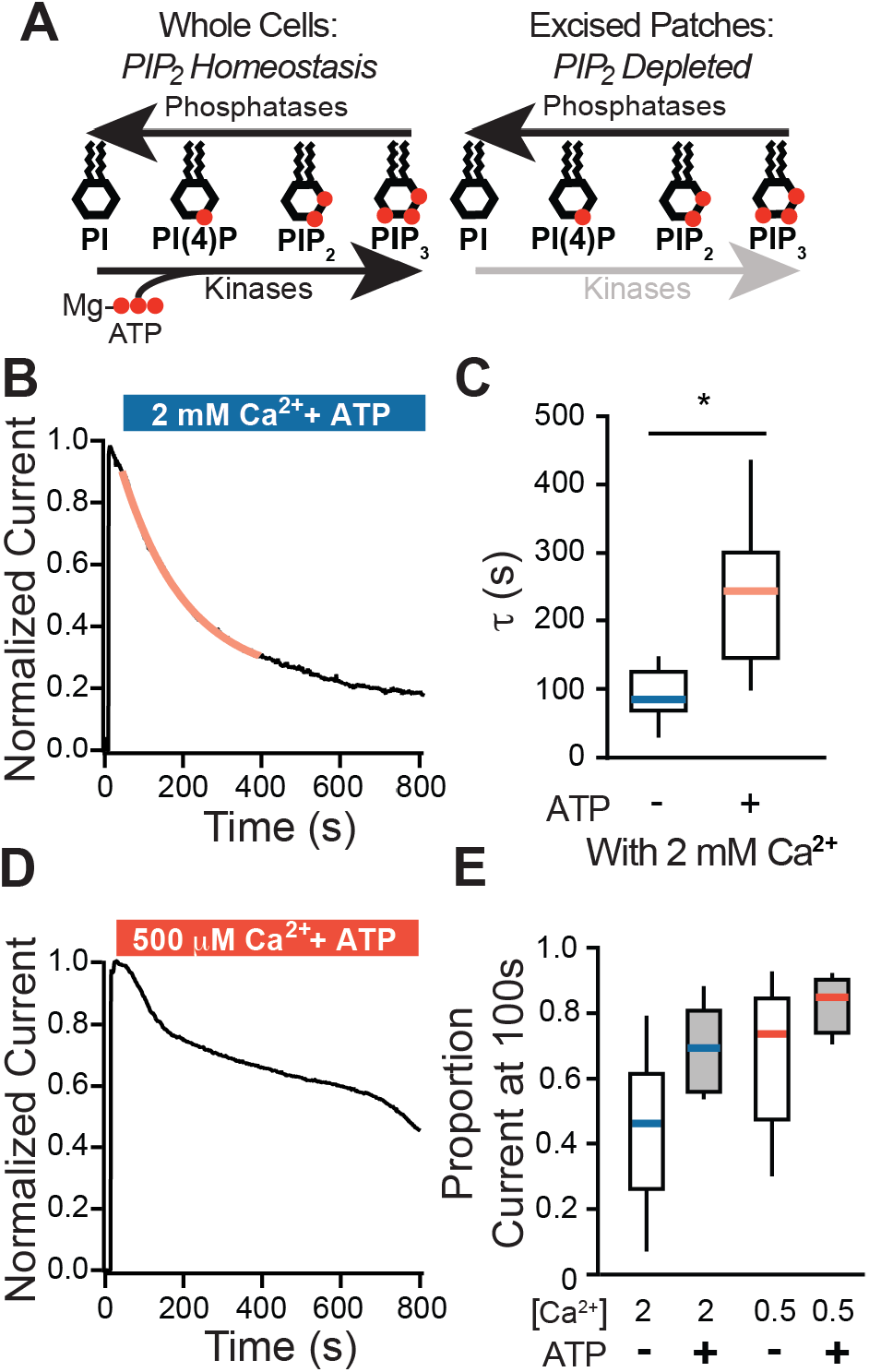
ATP application slowed current decay in excised inside-out patches. (A) Schematic illustrating the roles of phosphatases and kinases in maintaining phosphoinositide homeostasis in whole cells (*left*) vs excised patches (*right*). (B) Plot of normalized currents vs time during application of 2 mM Ca^2+^ with 1.5 mM Mg-ATP recorded from inside-out patches and fit with a single exponential (red line). (C) Box plot distribution of rate of current decay (τ) as reported by single exponentials, from patches treated with 2 mM Ca^2+^ with ATP or without ATP. (D) Example plot of TMEM16A conducted current before and during application of 1.5 mM Mg-ATP with 500 µM Ca^2+^ represented by the orange bar. (E) Box plot distribution of the proportion current at 100s in patches treated with indicated concentrations of Ca^2+^ (in mM) and with or without 1.5 mM Mg-ATP, as indicated (N=7-18 patches).

By fitting plots of the normalized current vs time with single exponentials, we observed that ATP application substantially slowed the τ of rundown from an average of 91.0 ± 8.3 s (N=18) in 2 mM Ca^2+^ alone, to 239.8 ± 13.6 s with ATP (N=7) (*P*<0.01, Figure 4C&D). A slowing of current decay was also observed in the proportion of maximum current at 100 s, from 0.45 ± 0.05 in 2 mM Ca^2+^ to 0.70 ± 0.05 with ATP (N=7-18). ATP also slowed current decay in 500 µM Ca^2+^; however, plots of current decay with 500 µM Ca^2+^ vs time were not well fit by single exponentials (Figure 4D). Again, we assayed for changes in current decay by quantifying the proportion of maximum current at 100 s, we observed a change from 0.67 ± 0.06 to 0.83 ± 0.03 (*P*=0.03, T-test, N=8-11). ATP slowed rundown with both 500 µM and 2 mM Ca^2+^, but rundown was still quicker in 2 mM Ca^2+^ with ATP. Together, these data suggest that millimolar Ca^2+^ speeds PI(4,5)P_2_ depletion by potentiating a process not recovered by ATP.

### 3.5 Inhibiting PLC slowed rundown of TMEM16A-conducted currents

We speculated that Ca^2+^ may activate PLC to thereby increase PIP_2_ cleavage for the generation of IP_3_. Importantly, the PI(4,5)P_2_ lost due to PLC cleavage cannot be recovered with ATP application. We verified that PLCs are present in the *X. laevis* eggs using a proteome dataset from fertilization-competent eggs (Wuhr et al., 2014) and queried for all proteins encoded by known PLC genes. Notably, this dataset was acquired on fertilization-competent eggs, but we are using immature stave VI oocytes for our experiments. Therefore, we also examined an RNA-sequencing dataset acquired from different stages of development of *X. laevis* oocytes and eggs (Session et al., 2016). We reasoned that if a protein is present in the stage VI oocyte, RNA should be present in the developing oocyte. Indeed, two PLC isoforms fit these criteria: PLCγ1 (encoded by the PLCG1 gene) and PLCβ3 (encoded the PLCB3 gene) (Supplemental figure 1).

We next determined whether inhibition of PLCs slowed rundown of TMEM16A currents. To do so, we used two PLC inhibitors: U73122 (Jin, Lo, Loh & Thayer, 1994) and PMSF (Walenga, Vanderhoek & Feinstein, 1980). We found that TMEM16A conducted currents ran down more slowly in the presence of 1 µM U73122 compared to application of 2 mM Ca^2+^ alone. In 1 µM U73122, the proportion of remaining current at 100 s was 0.71 ± 0.08 (N=5), compared to 0.45 ± 0.05 (N=18) in Ca^2+^ alone (*P*= 0.02, T-test). 2 mM PMSF similarly slowed rundown, with 0.78 ± 0.05 (N=6) proportion of remaining current at 100 s. Together, these data demonstrate that in addition to activating the channel, 2 mM Ca^2+^ also activates PLC to speed the rundown of TMEM16A conducted current.

## Discussion & Conclusions

Here we demonstrate that in addition to activating TMEM16A channels, intracellular Ca^2+^ speeds current rundown by turning on membrane tethered PLCs in excised inside out patches. TMEM16A channels require both intracellular Ca^2+^ and PIP2 to open and conduct Cl^−^ currents (De Jesus-Perez et al., 2017; Le, Jia, Chen & Yang, 2019; Ta, Acheson, Rorsman, Jongkind & Tammaro, 2017; Tembo et al., 2022; Tembo, Wozniak, Bainbridge & Carlson, 2019; Yu, Jiang, Cui, Tajkhorshid & Hartzell, 2019). Like other channels regulated by PIP_2_, TMEM16A-conducted currents decay when recorded in the inside-out configuration of the patch clamp technique (Figure 1), even in the continued presence of Ca^2+^ (De Jesus-Perez et al., 2017; Tembo, Wozniak, Bainbridge & Carlson, 2019). We previously demonstrated that this current decay from excised patches can be sped by scavenging membrane PIP_2_, slowed by inhibiting the membrane associated phosphatases or refueling the kinases (Tembo, Wozniak, Bainbridge & Carlson, 2019).

Our finding that TMEM16A-conducted currents decayed more slowly when the channels were activated by the other divalent cations Ni^2+^ or Ba^2+^ (Figure 2) provided our first hint that Ca^2+^ may target the channel and independently regulate PIP_2_ present in excised patches. This hypothesis was further supported by the demonstration that TMEM16A conducted current rundown was slower with application of a lower concentration of Ca^2+^, 500 µM (Figure 3).

Furthermore, ATP application slowed rundown in both 2 mM and 500 µM Ca^2+^, however, rundown was slower and more current remained when ATP was applied with 500 µM Ca^2+^ compared to 2 mM, indicating that another process contributed to rundown in higher Ca^2+^ (Figure 4).

PLCs can be activated directly by Ca^2+^ (Hwang, Oh, Shin, Kim, Ryu & Suh, 2005; Ryu, Suh, Cho, Lee & Rhee, 1987), we therefore speculated that Ca^2+^ may activate a membrane-associated PLC to speed rundown. Here we used bioinformatics to verify that indeed PLCs are present in the developing gamete: PLCγ1 and PLCβ3 (Supplemental Figure 1). To more directly test whether PLC depletion of PIP2 speeds TMEM16A current decay in excised patches, we quantified the rate of rundown with application of two different PLC inhibitors: U73122 and PMSF. Both inhibitors significantly slowed rundown in 2 mM Ca^2+^, thereby suggesting that PLC activity indeed contributed to loss of TMEM16A conducted current.

Here we report plots of current decay versus time using varying conditions, some of which were fit by single exponentials and others which were not. For example, plots of rundown in 2 mM Ni^2+^ were not well fit by single exponentials whereas plots in 2 mM Ba^2+^ were. The mechanism underlying this variability is not yet clear. Theoretically, fitting plots of relative current versus time with a single exponential would indicate that a single process gives rise to rundown. However, our data suggest that at least in 2 mM Ca^2+^, activity of both phosphatase and PLC deplete patch PI(4,5)P_2_. Further work is needed to understand this variability.

Fertilization activates TMEM16A channels in *X. laevis* eggs to allow Cl^−^ to leave the cytoplasm and depolarize the membrane (Wozniak, Phelps, Tembo, Lee & Carlson, 2018). This fertilization-signaled depolarization rise rapidly stops multiple sperm from entering an already fertilized egg, a process known as the fast block to polyspermy (Wozniak & Carlson, 2020).

Indeed, sperm can bind but will not enter depolarized eggs (Jaffe, 1976). Fertilization activates TMEM16A in *X. laevis* by a pathway requiring PLC activation and IP_3_ signaled Ca^2+^ release from the ER (Wozniak, Tembo, Phelps, Lee & Carlson, 2018) and occurs within seconds of fertilization. Fertilization also initiates a slower block to polyspermy that also require PLC and Ca^2+^ release from the ER to signal exocytosis of cortical granules from the egg minutes after sperm entry. The Ca^2+^ wave that initiates cortical granule exocytosis starts at the site of sperm entry and takes minutes to traverse the egg (Busa & Nuccitelli, 1985; Fontanilla & Nuccitelli, 1998). Whether the same process activates the PLC needed for the fast and slow polyspermy blocks in *X. laevis* eggs has not yet been determined.

A prominent role for Ca^2+^ in the regulation of PIP_2_ and TMEM16A is intriguing because both Ca^2+^ and PIP_2_ are required for the channel to open, yet elevated Ca^2+^ could signal deactivation of the channel even when Ca^2+^ is bound to the channel. Although here we report that millimolar Ca^2+^ concentrations were required to activate PLC and speed rundown, *X. laevis* eggs experience prolonged periods of high intracellular Ca^2+^ shortly after fertilization (Wagner et al., 2004). It is possible that this Ca^2+^ mediated PLC activation could give rise to the regenerative Ca^2+^ wave that initiates *X. laevis* cortical granule exocytosis (Fall, Wagner, Loew & Nuccitelli, 2004; Wagner et al., 2004). For example, fertilization could produce an acute and localized bolus of Ca^2+^ in *X. laevis* eggs. This Ca^2+^ could then activate nearby PLCs, which would then cleave PIP_2_ to generate IP_3_ and signal the release of more Ca^2+^. This mechanism would give rise to a self-sustaining and regenerating Ca^2+^ wave (Lechleiter, Girard, Peralta & Clapham, 1991; Lechleiter & Clapham, 1992; Marchant, Callamaras & Parker, 1999).

## Acknowledgments

We thank G.V. Hammond for helpful discussions and advice and D. Summerville for excellent technical assistance. This work was supported by an American Heart Association Predoctoral Fellowship 18PRE33960391 to M.T. and National Institute for Health grant 1R01GM125638 to A.E.C.

## Conflict of Interest

The authors declare no conflicts of interest in the contents of this manuscript.

**Figure 5:**
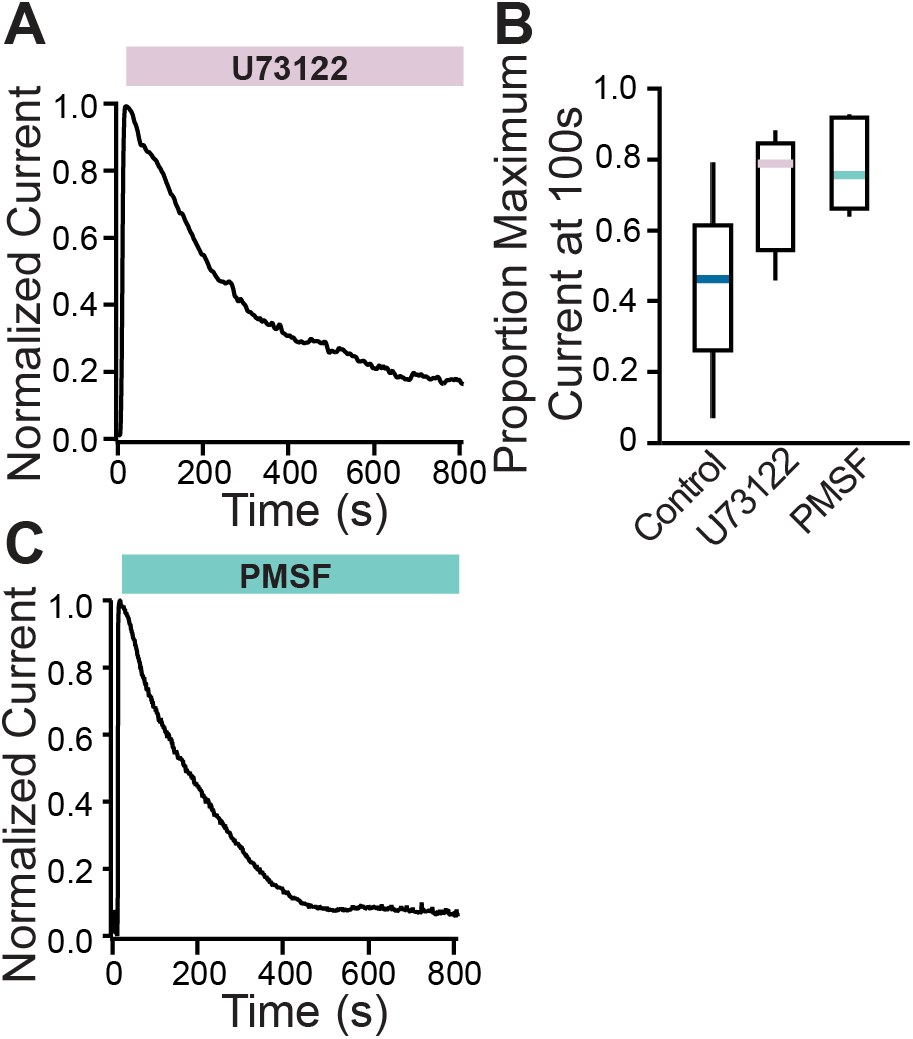
Inhibiting PLC slowed TMEM16A rundown. Representative plots of normalized current recorded at -60 mV vs time of TMEM16A conducted currents in the presence of 2 mM Ca^2+^ with 1 µM U73122 (A) or 2 mM PMSF (C). (B) Box plot distribution of rate of current decay (τ) as reported by single exponentials, from patches treated with 2 mM Ca^2+^ alone (control) or with U73122 or PMSF, as indicated (N=5-18).

**Figure 6:**
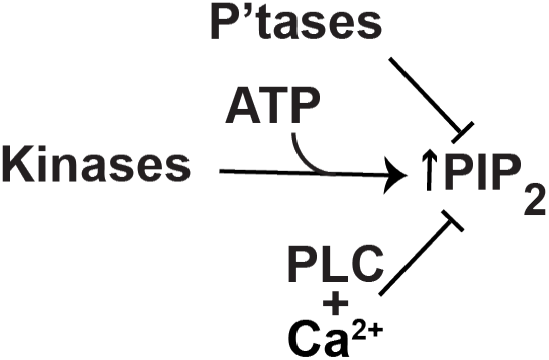
Ca^2+^ regulates membrane PI(4,5)P_2_. In excised patches, PI(4,5)P_2_ levels are regulated by membrane anchored enzymes including phosphatases, kinases, and PLCs.

## References

Al-Hosni R, Ilkan Z, Agostinelli E, & Tammaro P (2022). The pharmacology of the TMEM16A channel: therapeutic opportunities. Trends Pharmacol Sci 43: 712–725.

Bers DM, Patton CW, & Nuccitelli R (2010). A practical guide to the preparation of Ca(2+) buffers. Methods Cell Biol 99: 1–26.

Brunner JD, Lim NK, Schenck S, Duerst A, & Dutzler R (2014). X-ray structure of a calcium-activated TMEM16 lipid scramblase. Nature 516: 207–212.

Bulley S, Neeb ZP, Burris SK, Bannister JP, Thomas-Gatewood CM, Jangsangthong W, et al. (2012). TMEM16A/ANO1 channels contribute to the myogenic response in cerebral arteries. Circ Res 111: 1027–1036.

Busa WB, & Nuccitelli R (1985). An elevated free cytosolic Ca^2+^ wave follows fertilization in eggs of the frog, Xenopus laevis. J Cell Biol 100: 1325–1329.

Cabrita I, Benedetto R, Wanitchakool P, Lerias J, Centeio R, Ousingsawat J, et al. (2021). TMEM16A Mediates Mucus Production in Human Airway Epithelial Cells. Am J Respir Cell Mol Biol 64: 50–58.

Chen Q, Kong L, Xu Z, Cao N, Tang X, Gao R, et al. (2021). The Role of TMEM16A/ERK/NK-1 Signaling in Dorsal Root Ganglia Neurons in the Development of Neuropathic Pain Induced by Spared Nerve Injury (SNI). Mol Neurobiol 58: 5772–5789.

Dang S, Feng S, Tien J, Peters CJ, Bulkley D, Lolicato M, et al. (2017). Cryo-EM structures of the TMEM16A calcium-activated chloride channel. Nature 552: 426–429.

Danielsson J, Kuforiji AS, Yocum GT, Zhang Y, Xu D, Gallos G, et al. (2020). Agonism of the TMEM16A calcium-activated chloride channel modulates airway smooth muscle tone. Am J Physiol Lung Cell Mol Physiol 318: L287–L295.

De Jesus-Perez JJ, Cruz-Rangel S, Espino-Saldana AE, Martinez-Torres A, Qu Z, Hartzell HC, et al. (2017). Phosphatidylinositol 4,5-bisphosphate, cholesterol, and fatty acids modulate the calcium-activated chloride channel TMEM16A (ANO1). Biochim Biophys Acta 1863: 299–312.

Fall CP, Wagner JM, Loew LM, & Nuccitelli R (2004). Cortically restricted production of IP3 leads to propagation of the fertilization Ca2+ wave along the cell surface in a model of the Xenopus egg. J Theor Biol 231: 487–496.

Fontanilla RA, & Nuccitelli R (1998). Characterization of the sperm-induced calcium wave in Xenopus eggs using confocal microscopy. Biophys J 75: 2079–2087.

Hawn MB, Akin E, Hartzell HC, Greenwood IA, & Leblanc N (2021). Molecular mechanisms of activation and regulation of ANO1-Encoded Ca(2+)-Activated Cl(-) channels. Channels (Austin) 15: 569–603.

Hernandez-Clavijo A, Sarno N, Gonzalez-Velandia KY, Degen R, Fleck D, Rock JR, et al. (2021). TMEM16A and TMEM16B Modulate Pheromone-Evoked Action Potential Firing in Mouse Vomeronasal Sensory Neurons. eNeuro 8.

Hilgemann DW, & Ball R (1996). Regulation of cardiac Na^+,Ca2+^ exchange and KATP potassium channels by PIP2. Science 273: 956–959.

Huang F, Zhang H, Wu M, Yang H, Kudo M, Peters CJ, et al. (2012). Calcium-activated chloride channel TMEM16A modulates mucin secretion and airway smooth muscle contraction. Proc Natl Acad Sci U S A 109: 16354–16359.

Hwang JI, Oh YS, Shin KJ, Kim H, Ryu SH, & Suh PG (2005). Molecular cloning and characterization of a novel phospholipase C, PLC-eta. Biochem J 389: 181–186.

Jaffe LA (1976). Fast block to polyspermy in sea urchin eggs is electrically mediated. Nature 261: 68–71.

Jin W, Lo TM, Loh HH, & Thayer SA (1994). U73122 inhibits phospholipase C-dependent calcium mobilization in neuronal cells. Brain Res 642: 237–243.

Ko W, & Suh BC (2021). Differential Regulation of Ca(2+)-Activated Cl(-) Channel TMEM16A Splice Variants by Membrane PI(4,5)P2. Int J Mol Sci 22.

Le SC, Jia Z, Chen J, & Yang H (2019). Molecular basis of PIP2-dependent regulation of the Ca(2+)-activated chloride channel TMEM16A. Nat Commun 10: 3769.

Lechleiter J, Girard S, Peralta E, & Clapham D (1991). Spiral calcium wave propagation and annihilation in Xenopus laevis oocytes. Science 252: 123–126.

Lechleiter JD, & Clapham DE (1992). Molecular mechanisms of intracellular calcium excitability in X. laevis oocytes. Cell 69: 283–294.

Marchant J, Callamaras N, & Parker I (1999). Initiation of IP(3)-mediated Ca(2+) waves in Xenopus oocytes. EMBO J 18: 5285–5299.

Paulino C, Kalienkova V, Lam AKM, Neldner Y, & Dutzler R (2017). Activation mechanism of the calcium-activated chloride channel TMEM16A revealed by cryo-EM. Nature 552: 421–425.

Peters CJ, Gilchrist JM, Tien J, Bethel NP, Qi L, Chen T, et al. (2018). The Sixth Transmembrane Segment Is a Major Gating Component of the TMEM16A Calcium-Activated Chloride Channel. Neuron.

Ryu SH, Suh PG, Cho KS, Lee KY, & Rhee SG (1987). Bovine brain cytosol contains three immunologically distinct forms of inositolphospholipid-specific phospholipase C. Proc Natl Acad Sci U S A 84: 6649–6653.

Session AM, Uno Y, Kwon T, Chapman JA, Toyoda A, Takahashi S, et al. (2016). Genome evolution in the allotetraploid frog Xenopus laevis. Nature 538: 336–343.

Shi S, Pang C, Guo S, Chen Y, Ma B, Qu C, et al. (2020). Recent progress in structural studies on TMEM16A channel. Comput Struct Biotechnol J 18: 714–722.

Suh BC, & Hille B (2008). PIP2 is a necessary cofactor for ion channel function: how and why? Annu Rev Biophys 37: 175–195.

Ta CM, Acheson KE, Rorsman NJG, Jongkind RC, & Tammaro P (2017). Contrasting effects of phosphatidylinositol 4,5-bisphosphate on cloned TMEM16A and TMEM16B channels. British journal of pharmacology 174: 2984–2999.

Tembo M, Bainbridge RE, Lara-Santos C, Komondor KM, Daskivich GJ, Durrant JD, et al. (2022). Phosphate position is key in mediating transmembrane ion channel TMEM16A-phosphatidylinositol 4,5-bisphosphate interaction. J Biol Chem 298: 102264.

Tembo M, Wozniak KL, Bainbridge RE, & Carlson AE (2019). Phosphatidylinositol 4,5-bisphosphate (PIP2) and Ca(2+) are both required to open the Cl(-) channel TMEM16A. J Biol Chem 294: 12556–12564.

Wagner J, Fall CP, Hong F, Sims CE, Allbritton NL, Fontanilla RA, et al. (2004). A wave of IP3 production accompanies the fertilization Ca2+ wave in the egg of the frog, Xenopus laevis: theoretical and experimental support. Cell Calcium 35: 433–447.

Walenga R, Vanderhoek JY, & Feinstein MB (1980). Serine esterase inhibitors block stimulus-induced mobilization of arachidonic acid and phosphatidylinositide-specific phospholipase C activity in platelets. J Biol Chem 255: 6024–6027.

Wang H, Ma D, Zhu X, Liu P, Li S, Yu B, et al. (2021). Nimodipine inhibits intestinal and aortic smooth muscle contraction by regulating Ca(2+)-activated Cl(-) channels. Toxicol Appl Pharmacol 421: 115543.

Wozniak KL, & Carlson AE (2020). Ion channels and signaling pathways used in the fast polyspermy block. Mol Reprod Dev 87: 350–357.

Wozniak KL, Phelps WA, Tembo M, Lee MT, & Carlson AE (2018). The TMEM16A channel mediates the fast polyspermy block in Xenopus laevis. J Gen Physiol.

Wozniak KL, Tembo M, Phelps WA, Lee MT, & Carlson AE (2018). PLC and IP3-evoked Ca(2+) release initiate the fast block to polyspermy in Xenopus laevis eggs. J Gen Physiol.

Wuhr M, Freeman RM, Jr., Presler M, Horb ME, Peshkin L, Gygi SP, et al. (2014). Deep proteomics of the Xenopus laevis egg using an mRNA-derived reference database. Curr Biol 24: 1467–1475.

Yu K, Jiang T, Cui Y, Tajkhorshid E, & Hartzell HC (2019). A network of phosphatidylinositol 4,5-bisphosphate binding sites regulates gating of the Ca(2+)-activated Cl(-) channel ANO1 (TMEM16A). Proc Natl Acad Sci U S A 116: 19952–19962.

Yuan H, Gao C, Chen Y, Jia M, Geng J, Zhang H, et al. (2013). Divalent cations modulate TMEM16A calcium-activated chloride channels by a common mechanism. J Membr Biol 246: 893–902..

