## Supplemental Figure 1 for "PIP_2_ and Ca^2+^ regulation of TMEM16A currents in excised inside-out patches"

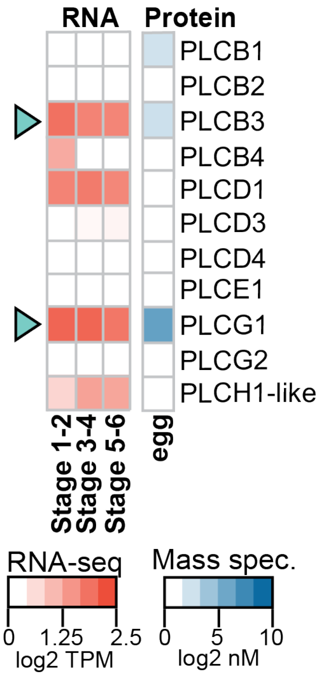


**Supplemental Figure 1: Expression of PLCs in *X. laevis* oocytes and eggs.**
Heatmaps of expression levels of PLCs at the developmental stages indicated. *Left*: Transcript levels (shown as transcripts per million [TPM]; from (Session et al., 2016)) RNA expression as determined by RNA-seq-based transcriptome study. *Right*: Protein concentrations (from Wuhr et al., 2014) as determined by mass spectrometry-based proteomics study (in $log_2$nanomolar). Arrowheads highlight PLCs present.
